# Global Quantitative Analysis of Ligation Reactions in Self-Assembled DNA Nanostructures at the Single-Nick Level

**DOI:** 10.1101/2025.06.18.660358

**Authors:** Konrad Hacker, Emilia Juricke, Carolin Münch, Antonio Suma, Adrian Keller, Yixin Zhang

**Author notes:** These authors contribute equally to this work (Konrad Hacker, Emilia Juricke). For correspondence: A.S.; A.K.; Y.Z.

## Abstract

Ligation of staple strands in DNA origami nanostructures (DONs) can yield enhanced structural stability in critical environments. This process can be viewed as performing hundreds of parallel reactions programmed on a self-assembled nanoscale platform. While previous studies have focused on investigating the collective results of the chemical or enzymatic ligation reactions, herein, the global quantitative analysis of individual ligation reactions is achieved using quantitative PCR (qPCR). By mapping enzymatic ligation efficiency on a trapezoidal substructure representing one third of a triangular DON, ligation is shown to preferentially occur at the trapezoid edges rather than at inner sites. Excellent agreement between the experimental ligation yields and docking simulations suggests that this is a result of variations in the ligase docking probability. Ligation products involving more than two consecutive sequences can be generated with each enzyme-catalyzed reaction as an independent event. Interestingly, the sharp contrast between the edges vs the inner sites has been abolished by changing the reaction condition and performing the ligation in a DMSO co-solvent system. This analytic method provides unprecedented insight into the multiple ligation reactions occurring in parallel within complex DONs and will be an invaluable tool in the translation of DONs from the lab to real-world applications.

## Introduction

Conventional chemical reactions are performed by mixing certain reactants to produce a desired product, and high reactant concentrations are favorable to drive the reaction to completion with high rate. In biology, while many chemical reactions are taking place simultaneously, nature controls numerous reactions by using the proximity effect to modulate the effective molarity of highly dilute reactants with macromolecules as templates. By mimicking nature’s approach, DNA-templated organic synthesis (DTS) has emerged as a general technique to control the reactivity of synthetic molecules by bringing them into proximity through attaching them to two DNA sequences complimentary to each other [1] (figure **1A**). DTS has already found many applications in constructing DNA-encoded chemical libraries [2]. However, their designs remain much simpler than many highly orchestrated biological processes such as protein syntheses on ribosomes, as DTS reactions are mostly governed by the recognition and Watson-Crick pairing between two freely diffusing DNA sequences. Templates with more advanced architectures would be needed to realize more sophisticated reactions involving multiple components, ideally with delicate arrangements in 2D or 3D to spatially program the reactions.

**Figure 1.**
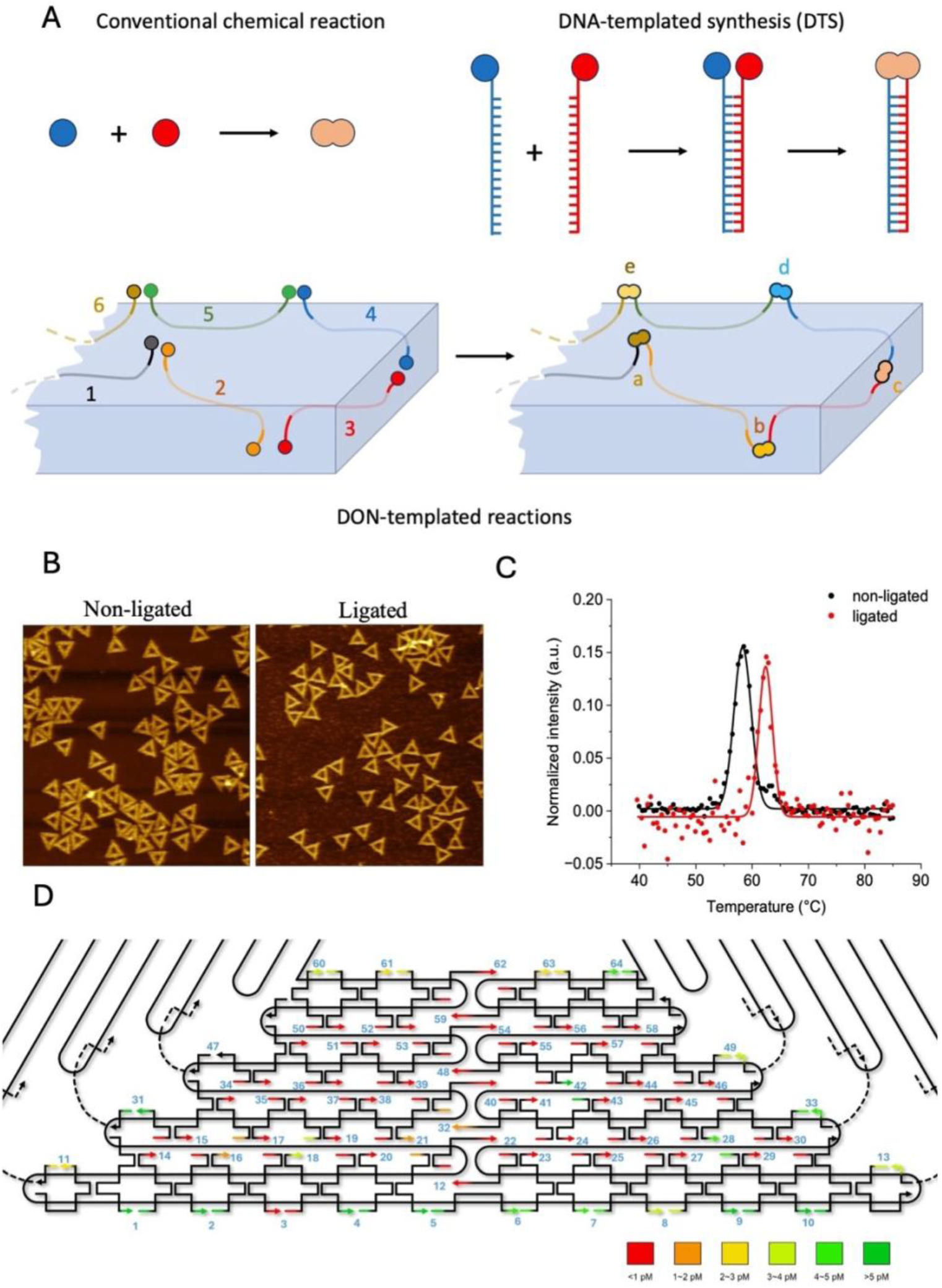
*Quantification of DON-templated chemical reaction.* (**A**) Control of the reactivity between two chemical groups in a free-diffusion system, as in conventional chemical reactions; or in a DNA-templated form; or templated by DNA origami nanostructure (DON). (**B**) AFM images (1.5 × 1.5 µm^2^) of triangular DNA origami structures with and without enzymatic ligation by T4 DNA ligase. Height scales for both images are 3.6 nm. (**C**) Melting curves of triangular DNA origami structures with and without enzymatic ligation by T4 DNA ligase. (**D**) Mapping of 64 ligation reactions on DON, in an entire trapezoidal domain, and in the regions between the domains. The ligation yields are presented as color-coded heatmap.

Looking for more complex architectures that can serve as templates for chemical reactions, we settled on self-assembled DNA origami nanostructures (DONs) as a particularly promising approach [3, 4]. By folding a long single-stranded viral DNA by hybridization with many short staple strands, the DNA origami technique enables the versatile construction of materials over scales from nanometers to micrometers with sub-nm addressability [5]. It is thus utilized for various applications in materials science [6], optics [7], robotics [8], biology [9], and medicine [10]. In many of these applications, DONs are exposed to destabilizing environments, resulting in shape distortions, melting, and degradation [11, 12, 13]. Consequently, different methods have been reported to improve DON stability through conjugating neighboring staple strands chemically or enzymatically. For example, Ramakrishnan et al. have used T4 DNA ligase to ligate the staple strands in triangular DONs, resulting in improved stability under denaturing conditions [14]. Enzyme-free ligation strategies have been reported by using cyanogen bromide (CNBr) [15] or 1-[(3-dimethylamino)propyl]-3-ethylcar-bodiimide (EDC) [16], to form phosphodiester bonds between staple strands through chemical condensation. While hundreds of ligation reactions take place simultaneously in the DONs, gel analyses have shown that such systems are highly heterogeneous. A quantitative analysis that evaluates each ligation reaction individually will shed new light on the contribution of ligation reaction conditions as well as ligation efficiency at individual nick sites to overall DON stability and thus provide an invaluable tool for assessing and optimizing DON ligation efficiency to enable their translation to real-world applications.

Apart from their practical application in stabilizing DONs in challenging environments, such reaction systems also offer advanced architectures for studying DNA-templated chemical reactions with sophisticated spatial arrangements in 2D and 3D. While conventional DTS studies can only investigate reactions between two compounds, the DNA strands arranged in a 2D or 3D template will allow reactions involving multiple components. Will reaction *A* ligating sequences *1* and *2* with a yield of *a*, and reaction *B* ligating sequences *2* and *3* with a yield of *b* occur independently of each other (yield of ligation product *1-2-3 Y* = *a×b*)? Or are the two reactions agonistic (*Y* > *a×b*) or antagonistic (*Y* < *a×b*) to each other? Such studies could result in artificial systems recapitulating the complexity and responsiveness of biology. However, this will be possible only if we have a robust analytic tool to quantify all intermediates and products. As the difficulty of studying a biological system quantitatively increases exponentially with the number of components involved, carrying out such analyses with a complex system on a global level will allow us to develop models for quantitative biology. For example, ligating all 208 staple stands in a Rothemund triangle [17] can be viewed as > 200 templated reactions, though the DON was not originally designed for this purpose. If ligation efficiency were the only factor, this would result in tens of thousands of possible products of varied length, while non-templated reactions would generate an infinite number of variations.

In this work, we have used quantitative PCR (qPCR) to perform a global analysis of enzyme-catalyzed ligation reactions in a triangular DON (Scheme **S1**). The primer pairs were meticulously designed to be specific for each ligation product. This allowed us to map the ligation reaction yields on one third of the whole DON and provided us with insights into the effects of steric and dynamic environments on enzymatic ligation by comparing experimentally determined ligation yields to molecular docking simulations. While we have focused on the ligation between two strands in one trapezoidal domain of the DON triangle, the method also allowed us to investigate more sophisticated reactions involving more than two components, to address the questions whether the reactions are independent from each other, agonistic or antagonistic.

## Results and Discussion

### Effect of Enzymatic Ligation on DON Stability

As can be seen in the atomic force microscopy (AFM) images in figure 1B, enzymatic ligation does not result in notable changes in the overall DON shape. Nevertheless, the melting transition of the ligated DON triangle is shifted to higher temperature (figure 1C and S1). Under Mg^2+^-free conditions, the melting temperature was found to increase from 58.3 to 62.4 °C upon ligation. While this increase is in line with previous reports [14, 15, 18]it appears rather moderate considering the large number of staple nicks in the triangular DON (208). In the absence of any additives such as DMSO, which was found to improve ligation reaction yields considerably [15], ligation efficiencies in 2D DONs typically range from 31 to 55%, depending on the reaction conditions [15, 18]. While the reason for these low ligation yields so far is unclear, previous works with nuclease enzymes suggest that certain DON design parameters may prevent the binding of the enzymes to specific locations in the DONs [19, 20, 21, 22]. Therefore, we set out to quantify the local ligation efficiency at individual nicks and evaluate how they are affected by the local DON environment.

### Quantification of DONs

To calculate the reaction yields, the concentrations of the Rothemund triangles used as starting materials first have to be quantified. However, quantification measurements based on UV absorption may suffer from experimental artefacts as they cannot distinguish the DONs from residual staples with higher absorbance and may be affected also by incomplete base pairing and scattering effects [23, 24]. As DNA origami are synthesized by mixing the DNA scaffold with a large excess of staple strands, we can quantify the scaffold DNA using qPCR with primer pairs specific to the scaffold DNA but not to any staple strands and ligation products. A calibration curve was obtained by performing qPCR measurements with a synthetic 64 nt DNA sequence from the scaffold in serial dilutions. The same primers were then used to quantify a sample of triangular DONs of 5.0 pM, a concentration estimated by UV absorption, resulting in a concentration of 5.8 pM (figure **S2**).

### Establishment of the Assay

Ten DNA sequences, corresponding to ten ligation products in one trapezoidal domain (figure 1), were randomly chosen to establish the quantification protocol by qPCR. Pairs of primers were designed, which are specific for every particular ligation product, amplifying neither any non-ligated staple strands, nor any other ligation products, nor the scaffold. A calibration curve was established for each DNA sequence using synthetic sequences in serial dilutions (figure **S3**). Each primer pair was then used to quantify the corresponding ligation product in ligated DONs, and compared to DONs not being subjected to enzymatic ligation as a negative control (figure **S4**). While the primers are not able to amplify the staples in non-ligated DONs, the qPCR quantifications revealed ligation reactions with yields in a broad range of < 1% to ∼ 100% (figure **S5**).

It is cumbersome and expensive to perform a calibration experiment for every ligation product. Therefore, we wondered, whether a single calibration curve could be used for all experiments. We have compared the calculated concentrations using either individual calibration curves or a universal calibration curve (e.g., the calibration curve for the ligation product of nick 1). Differences not larger than one qPCR cycle have been measured (figure **S5**). Therefore, for the following global quantification of ligation reactions in one trapezoidal domain of the triangular DON, the calibration curve for the ligation product of nick 1 was used to calculate the reaction yields.

### Global Assessment of Ligation Reactions

The ligated and non-ligated DONs were subjected to 64 qPCR quantification experiments, mapping the reactions not only in an entire trapezoidal domain, but also the regions between the domains (figure **1D**). It becomes apparent that most reactions with low performance can be found in the center of the trapezoid. On the contrary, reactions with high performance are observed mostly at the edges, especially at the outer edge and between the trapezoidal domains. There are also a few outliers, for example, the low yield of nick 3 at the outer edge and the high yield of nick 42 in the center.

We have taken a shotgun approach to demonstrate that our method can be used in large scale analysis without causing false positives. From the 64 forward primers and 64 backward primers (Table **S2**), 82 pairs have been randomly generated and used in the qPCR analysis. As shown in figure **S6**, none of the pairs can generate a clear qPCR signal. Therefore, ligation on DONs can take place only at the nick sites, while the shot gun experiment can exclude the possibility of producing high qPCR signals either from scaffold or due to unspecific ligation reactions among staple strands.

### Docking Probability from Molecular Dynamics Simulations

To understand the differences among the templated ligation reactions in the DON, we computed the probability to dock the T4 DNA ligase to each of the various nick sites, using a method developed in Ref. [21]. First, we started from conformations of the DON structures sampled using molecular dynamics simulations with the second version of oxDNA [25], a coarse-grained DNA model that has been shown to accurately reproduce the mechanical properties of double-stranded and single-stranded DNA molecules. These simulations, performed in Ref. [21], used Langevin dynamics and the software LAMMPS [26] [27]. Temperature was set at T = 300 K, and the monovalent salt concentration of the implicit solvent at 1 M. The timestep was chosen as 0.01τ_MD_, with τ_MD_ the simulation time unit, and simulations lasted about 3 · 10^7^τ_MD_, for a total of 3000 conformations saved. For each time frame, we superimposed through a rototranslation the crystal structure of the enzyme (PDB: 6DT1) to each nick and evaluated if the enzyme structure overlapped with the neighboring DNA strands. The percentage of DON conformations where the enzyme does not overlap with the DNA gives then the docking probability.

The simulations revealed three different types of docking sites (figure 2A).

**Figure 2.**
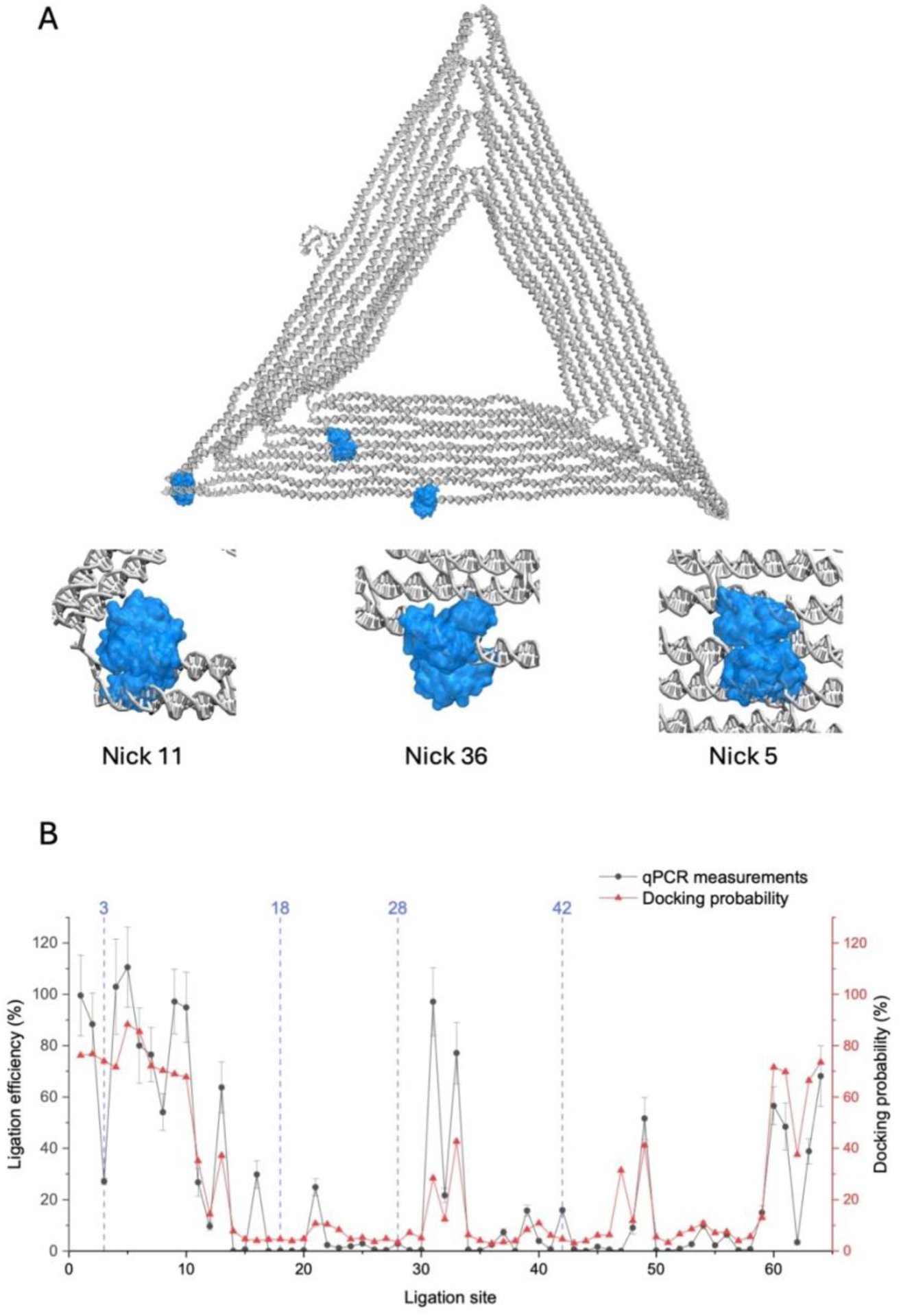
*Simulations of docking probability of T4 DNA ligase within the DON triangle.* (**A**) The crystal structure of the enzyme was rototranslated onto the DON structure that was sampled via oxDNA simulations. Nick 5 exemplifies the docking at the outer edge with high accessibility, while nick 36 exemplifies the docking within the trapezoid with low accessibility. Nick 11 represents the sites at the vertices and edges between the trapezoidal domains. (**B**) Comparison between the docking probabilities and the qPCR quantifications. While the qPCR measurements are highly reproducible in the technical replicates, the reaction yields at nicks 3, 18, 28, and 42 have been revised according to the batch-to-batch variation studies. Data are presented as mean values ± SD (n = 3).

The type I sites with the highest accessibility are located along the outer and inner edges of the trapezoid, such as nick 5.

The type II sites are the vertices and edges between the trapezoidal domains, such as nick 11. Although both neighboring domains can hinder the docking of DNA ligase, the gaps between domains restrict accessibility less dramatically than a continuous 2D surface.

The type III sites are located within the trapezoid, such nick 36. Due to the nature of planar structure, they exhibit the highest steric hinderance, resulting in the lowest docking probability of the ligase.

### Batch-to-batch variation analyses

By comparing the qPCR quantification experiments with the simulations (figure **2B**), there are multiple sites exhibiting remarkably higher yields in the qPCR quantification. Several factors might be responsible for these deviations. First, there may be differences in phosphorylation efficiency of the different staples. While T4 polynucleotide kinase (PNK) can phosphorylate oligonucleotides with any 5’-terminal nucleotides, it does show modest kinetic preferences, with single strands having a 5’ guanine being phosphorylated about six times more efficiently than single strands of similar length but featuring a 5′ cytosine [28]. These known differences, however, cannot account for the variations in ligation efficiency observed in our experiments. Consider for example nicks 31 and 42, which both showed a higher ligation efficiency than predicted by the docking simulations. However, the 5’ ends of the staples at both nick sites carry cytosines and thus should have lower phosphorylation efficiencies than the average staple. In contrast, nick 3 showed a markedly lower ligation efficiency than expected, even though the 5’ end of the staple carries a thymine. Second, simulations suggest that the DON triangles are not perfectly flat in solution but rather adopt a cup shape [21], which can affect the steric hinderance associated with a 2D surface. Third, the difference can also be caused by batch-to-batch variations. As the DONs are assembled from hundreds of staple strands through a kinetically and thermodynamically sophisticated process, some minimal change in experimental conditions may affect the resulting structure in an unpredictable manner. Moreover, subtle differences and defects caused by batch-to-batch variations cannot be revealed by standard gel electrophoresis or AFM analysis, in particular at the single-staple or single-nick level. As our qPCR analyses would allow us to investigate the presence and quantity of every ligation product, it can be used as a quality control method to reveal whether an outlier is just the result of batch-to-batch variation or intrinsic to the DNA-templated ligation reaction at a specific position.

We have first re-investigated the ligation at nick 42, as it is one of the most obvious outliers not in agreement with the simulation. We found the high yield is consistent in 5 technical replicates, each in triplicates (figure **3A**). Upon subjecting another independent DON sample preparation to qPCR analysis, a lower yield was measured, which is also consistent in 5 technical replicates, each in triplicates. Nevertheless, the ligation yield is still remarkably higher than those of all nick sites around it. We have then chosen 12 ligation reactions (figure **3B**), 8 in good agreement with the simulation with either high or low docking probability, and 4 not in agreement with the simulation, and compared them in four independent DON sample preparations. Interestingly, all 8 ligation reactions in good agreement with the simulation have exhibited consistent results, showing little batch-to-batch variations. For the three out of the four ligation reactions not in agreement with the simulation (figure **2B**), the three new preparations of DON samples have led to results in good agreement with the simulation (figure **3B**). The qPCR measurement, as a simple and quantitative assay, allowed us to reduce the mismatch between experiments and simulation by removing the artifacts resulted from batch-to-batch variations.

**Figure 3.**
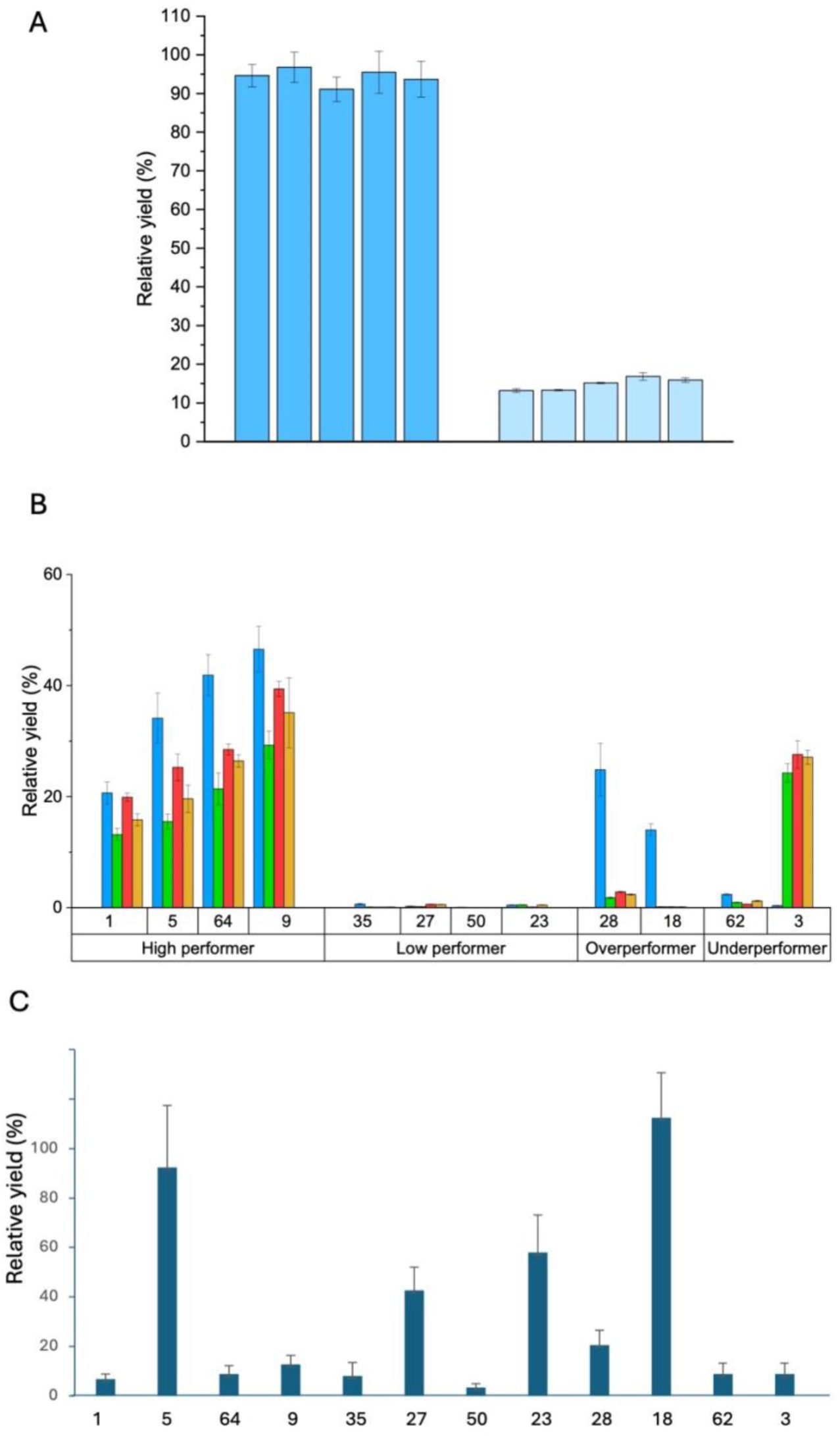
*Batch-to-batch variation and DMSO studies*. (**A**) Quantifications of the ligation at nick 42 with two independently prepared DON samples. Each sample was analyzed in 5 technical replicates, each in triplicates. (**B**) Quantifications of the ligation reactions at 12 nick sites with four independently prepared DON samples. Data are presented as mean values ± SD (n = 3). (**C)** Quantifications of the ligations at 12 nick sites with DONs ligated in the presence of DMSO. Data are presented as mean values ± SD (n = 6).

### Effect of DMSO on DON ligation

Krishnamurthy, Morii et al. [15] have reported a co-solvent DMSO-assisted enzymatic ligation method, resulting in more complete ligations as well as higher melting temperature. Indeed, adding DMSO to our ligation reaction resulted in a remarkably elevated melting temperature of 67.4 ± 0.3 °C (figure **S7**), as compared to a melting temperature of 58.4 ± 0.1 °C for DONs before the ligation, and a melting temperature of 62.6 ± 0.1 °C for DONs ligated in aqueous solution.

We have then quantified the ligations at the 12 nick sites, which had been chosen in the batch-to-batch variation study (figure **3B** and **3C**). Interestingly, different from the ligation reactions performed in aqueous solution, clear ligation products can be detected at all nick sites, with the lowest performers at nick positions of 1 and 50 having ligation yields of 3.4 ± 1.6 % and 6.6 ± 2.3 %, respectively. More remarkably, the pattern of edges *vs.* inner sites have been altered. At the edge, nick site 1 exhibited low ligation yield, whereas nick site 5 has shown a very high ligation yield of 92.3 ± 25 %. In contrast, at the inner sites, high ligation yields have been measured for nick sites 27, 23, and 18. As DMSO affects the mechanical properties of dsDNA [29] and thereby alters the dynamics of DONs, the coarse-grained molecular dynamics simulations established in aqueous solution can no longer be applied. The DMSO-assisted enzymatic ligation methodology can increase the DON thermal stability, not only through enhancing the ligation yields, but also through strengthen the inner nick sites, at which ligation yields are ubiquitously low under aqueous conditions.

### Local Fluctuations

While we have no available method to analyze or modulate local DON topology, local flexibility and dynamics of a selected nick can be enhanced by removing neighboring staple strands. For this, we selected nick 43, which has an extremely low ligation efficiency (figure **4A**). We investigated whether we could increase the ligation yield by removing the pair of staple strands either at its left side, at its right side, or at both sides. Interestingly, removing a single pair, either at the left side or at the right side, has little effect on the DON-templated reaction. Removing both pairs, however, resulted in a remarkably increased ligation yield. Therefore, subtle changes in the local environment of a nick site can affect its accessibility by the enzyme, and the steric hinderance associated with a continuous 2D surface can be mitigated by increasing the local fluctuations.

**Figure 4.**
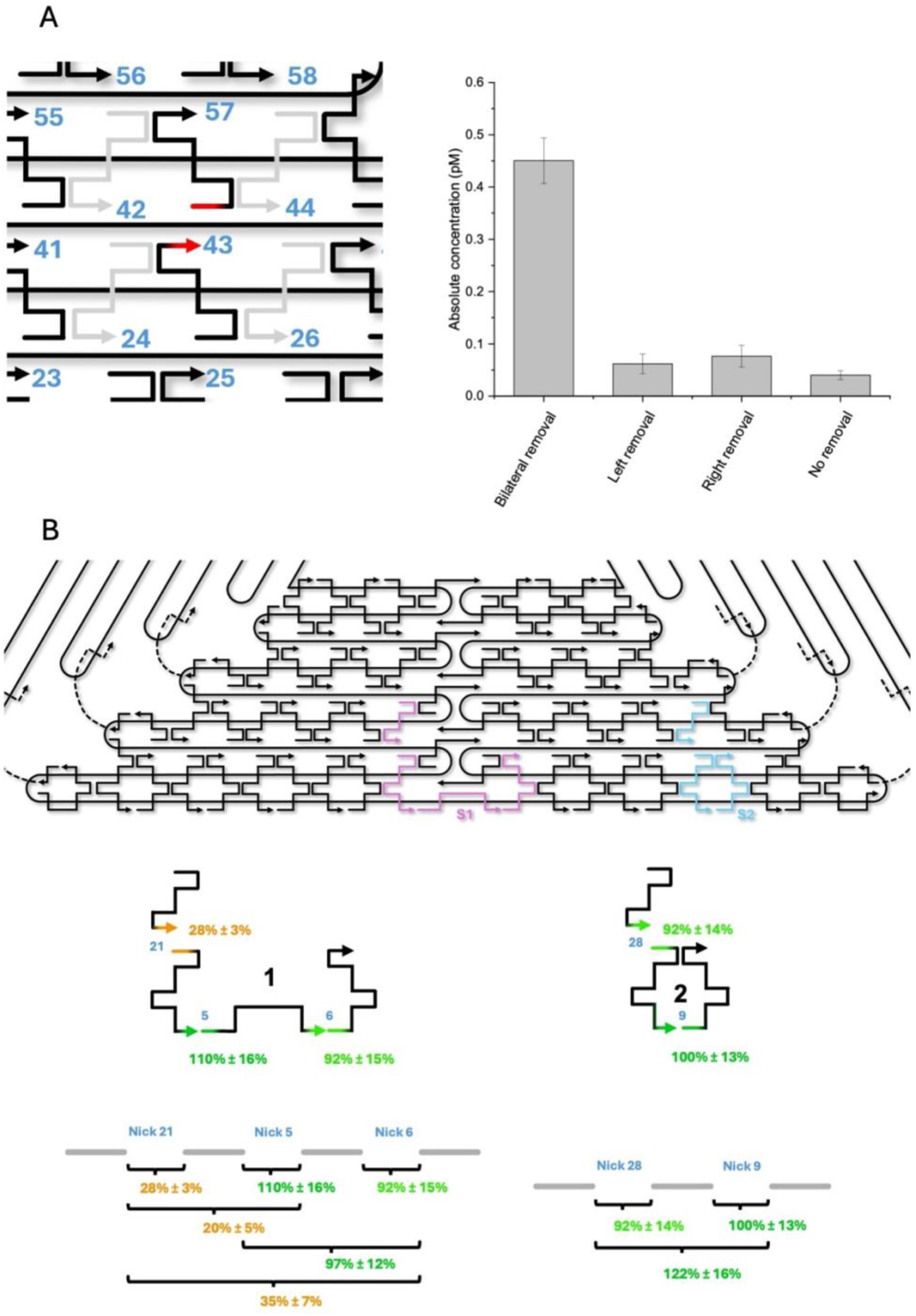
*Control of local dynamics and consecutive ligations*. (**A**) Increase of ligation yield at nick 43 by removing the neighbors. (**B**) Consecutive ligation of four DNA sequences on DON resulting in sequence 1 and consecutive ligation of three DNA sequences resulting in sequence 2. qPCR measurements can be used to quantify all intermediates and final products. Data are presented as mean values ± SD (n = 3).

We can also compare our results to the reactivity of a restriction endonuclease enzyme on the same DON structure, characterized experimentally in Ref. [19]and theoretically with the same docking method in Ref. [21]. In these works, it was shown that the action of the enzyme can be strongly affected by the position of the cleavage site, as well as through removal of selected staples. In particular, the endonuclease characterized presents a bulge, which can overlap with a DNA strand neighbor to the docked one. Based on the orientation of the GCGC sequence to be cleaved and its positioning inside the DON structure, one can have situations similar to the ligase, where the bulge does not overlap with other DNA strands (e.g. two type I sites), intermediate cases where overlaps happen around 50% of the time, and cases where overlap occurs all the time. However, we would like to stress that the ligase has two opposite bulges, which further restrict DNA binding, requiring enough space from two opposite directions. Moreover, the nick sites characterized here are always positioned at the same angle with respect to the DON surface, while in the case of the endonuclease the directionality is variable, rendering even type III sites completely cleavable if they have the correct angle.

### Consecutive Ligations

The qPCR method can quantify ligation products of only two staple strands, but also of multiple, consecutive staples. We have quantified the DNA sequence S1, which is the consecutive ligation product of four staple strands resulting from parallel ligation of nicks 21, 5, and 6, and analyzed its relationship with the three ligation reactions between two strands, as well as the two consecutive ligations among the three sequences. We have done the same also for the DNA sequence S2, which is the consecutive ligation product of three staple strands resulting from ligation of nick 28 and nick 9. As shown in figure 4B, the consecutive ligation products S1 and S2 could be successfully quantified. Remarkably, the yield of a consecutive ligation is approximately the multiplication product of individual ligation events. As neither strong agonistic nor antagonistic effects were observed, these results indicate that the ligation reactions catalyzed by the DNA ligase are independent events.

## Conclusion

DNA nanotechnology has become a highly interesting method for the development of therapeutic and diagnostic tools. However, as DONs are assembled from hundreds of staple strands through a kinetically and thermodynamically utterly sophisticated process, there is no reliable method for analyzing batch-to-batch variations. AFM can be used to evaluate their structural integrity but without resolving subtle differences and defects, while the precise 3D structures of such extremely complex assemblies in solution remain unknown. The qPCR-based method reported in this study can pave the way for the quantitative analysis of ligated DONs for their potential applications in biomedicine. Our experiments and simulations reveal the importance of the local environment surrounding a nick site for efficient ligation, which may be exploited on the DON design level to enhance ligation efficiency. Similar approaches already have resulted in DON designs with enhanced resistance toward nuclease degradation [22]. Moreover, as the reactions are programmable events on the self-assembled nanostructures, this analytic method could aid the development of new tools for genome assembly and DNA computation. It is important to note that qPCR analysis is most robust for single-stranded DNA with native bonds. PCR and qPCR involving non-native bonds or more advanced architectures [30] [31] will cause biases in the polymerization reaction by polymerase and thus be more suitable for qualitative and comparative analyses.

### Experimental Section

#### Phosphorylation of Staple Strands, DNA Origami Assembly, and Ligation

To enable ligation, the 5’ ends of all 208 staple strands (Eurofins) of the Rothemund triangle [17] were phosphorylated first using T4 PNK (New England Biolabs). A total of 1.35 nmol of staple strands (33.75 mM final concentration) were incubated with 45 U PNK in 1× PNK reaction buffer (New England Biolabs) supplemented with 1.5 mM ATP (New England Biolabs) in a total volume of 40 μL. The reaction mixture was incubated at 37 °C for 30 minutes, followed by thermal inactivation of the enzyme at 65 °C for 20 minutes.

Following phosphorylation, 10 nM scaffold p7249 (M13mp18, 100 nM stock, Tilibit) was added to the staple mixture. The folding solution was adjusted to 1× TAE (Roth) supplemented with 12.5 mM MgCl_2_ (Roth), yielding a final volume of 50 μL. Thermal annealing was performed using a Ristretto Thermocycler (VWR) by heating the solution to 80 °C, followed by gradual cooling to room temperature over 90 minutes. The cooling ramp consisted of an initial decrease to 55 °C at - 0.5 °C per 12 seconds, followed by a slower ramp to room temperature at -0.3 °C per 48 seconds.

DONs were purified using Amicon® Ultra-0.5 centrifugal filters with a 100 kDa molecular weight cutoff (Merck). Filters were pre-rinsed with 300 μL of 1× TAE containing 12.5 mM MgCl_2_ and centrifuged at 7000 g for 7 minutes. For purification, 50 to 100 μL of folded DON solution was diluted to 400 μL with the same buffer and centrifuged at 6000 g for 10 minutes. This was followed by an additional wash with 350 μL of buffer under identical conditions. DON triangles were recovered by inverting the filter und centrifuging at 7000 g for 7 minutes. The concentration of the purified structures was determined using a Nanophotometer P330 (Implen).

To compare ligated and non-ligated DONs, two samples were prepared from the same batch of phosphorylated structures: one with T4 DNA ligase (New England Biolabs) and one without. Ligation reactions were carried out in a total volume of 50 μL, containing 10 nM DON, 1× ligase reaction buffer and 1500 U of T4 DNA ligase. For DMSO-assisted ligation, 10 μL of DMSO (Merck) was added to achieve a final concentration of 20 % (v/v). Samples were incubated at 16 °C for 12 hours (overnight), then stored at 5 °C until purification. Protein removal and concentration determination were performed as described above, using HPLC-grade water (Roth) as the washing solvent [32].

#### qPCR Measurements

Each qPCR assay was conducted using Luna^®^ Universal qPCR Master Mix from New England Biolabs. This Master Mix contains SYBR Green I, a non-specific double-stranded DNA intercalating dye, along with other PCR reagents in unspecified amounts, to enable real-time measurement of PCR product formation. Each reaction was prepared with a total volume of 20 µL per well, in a 96-well microplate. The general assay consisted of 10 µL of Luna^®^ Universal Master Mix, 1 µL of a 5 µM premixed primer solution (final concentration 250 nM) in Milli-Q^®^ water, 8 µL of Milli-Q^®^ water and 1 µL of the target DNA. For consistency, aside from the target DNA, all components were premixed in the aforementioned ratios. Subsequently, 1 µL of target DNA was added to each well, followed by dispensing of 19 µL of the premixed solution.

To establish a calibration curve, an oligonucleotide matching the target sequence was prepared in tenfold serial dilutions from 1 nM to 0.1 pM. For quantification of ligation yield, each sample was prepared with 1 µL of a 120 pM DON sample stock solution quantified by scaffold amplification, resulting in a 6 pM final DON concentration, well within the established range of the calibration curve. The primers were designed and optimized using Primer3UT, a website specifically recommended by the manufacturers of the qPCR master mix (see supplementary information). The design focused on achieving an as low as possible difference in melting temperature, a high annealing temperature, a guanine-cytosine content of approximately 50%, and a low likelihood of forming homodimers, heterodimers, or hairpins. After each qPCR run, the microplate was cleaned with 70% ethanol and Milli-Q^®^ water and was, subsequently, left to dry overnight under the laboratory hood. Luna^®^ Universal Master Mix, primers, oligonucleotide standards, and DON samples were stored at -20°C. The respective aliquots were temporarily held at 4°C during sample preparation.

#### Thermal Cycler

Thermal cycler program as described in supplementary information was used. In general, reactions started with an initial denaturation phase at 95 °C for 1 minute to get rid of persistent secondary structures. After another 10 seconds at 95 °C, samples are cooled down to around 50 °C for 5 seconds for primer annealing, with the specific temperature being determined by the respective primer pair melting temperature. During subsequent extension at 72 °C for 5 seconds, the qTOWER performed the fluorescence signal measurement. After extension is completed, the three phases, 20 seconds in total, are repeated 40 times. Finally, a melting curve analysis with a ramping temperature from 60 °C to 95 °C over 8 minutes and 55 seconds was performed as qPCR product quality control.

#### Light Scattering

A JASCO fluorescence spectrometer FP-8200 with a water-cooled Peltier thermostatic cell holder was used to monitor the temperature-dependent light scattering. Before each measurement, the cuvette was rinsed with HPLC-grade water, followed by ethanol, and subsequently dried under a stream of argon. For each experiment, 350 μL of a 5 nM sample in HPLC-grade water was used. The intensity of scattered light at the maximum absorption of the DONs at 340 nm was recorded over time while applying a temperature gradient from 40 °C to 85 °C at a rate of 0.3 °C per minute. Melting temperatures were determined by applying a Gaussian fit to the first negative derivate of the normalized intensity curves.

#### AFM

100 μL of 1 nM of DON triangles in 1× TAE with 12.5 mM MgCl_2_ were adsorbed on freshly cleaved PELCRO® mica sheets (Ted Pella, Inc) for 5 minutes. Afterwards, the absorbed DONs were gently washed with 15 mL of HPLC-grade water and then dried under a stream of argon. AFM imaging was performed in air using a JPK Nanowizard Ultra Speed (JPK Instruments) in intermittent contact mode with HQ:NSC18/Al BS cantilevers (75 kHz, 2.8 N/m) from MikroMasch (NanoAndMore).

#### Statistical Analysis

A passive-reference dye is used to normalize the fluorescence signal. For this purpose, the raw fluorescence signal (RFU) of the intercalating SYBR^®^ Green I is divided by the RFU of a passive-reference dye at each cycle. The resulting value is subsequently offset by -1, yielding ΔR_n_. For each sample, a common baseline correction from cycle 1 to 5 was manually defined. For ΔR_n_ of each cycle of the run, the mean ΔR_n_ of the correction window was subtracted to set the starting values of amplification curves to zero. Using the instrument’s second-derivative maximum algorithm, the C_T_ value of a standard was set automatically. Amplification curves were smoothed with a moving average over 3 consecutive cycles with linear ΔR_n_ scaling. Samples were measured in triplicates, whereas two negative controls, non-ligated scaffold strand and NTC, were each measured once. Data of biological samples is presented as the geometric mean of the technical triplicates and the corresponding standard deviation in the format mean ± SD. The geometric mean was used to describe exponentially scaled data, as is the case with C_T_ values and their corresponding standard concentrations. According to MIQE guidelines, technical replicates with C_T_ values deviating by more than 0.5 cycles or exhibiting a standard deviation σ greater than 2 were repeated to ensure data quality. Relative yields necessitate an error propagation since ligation site and scaffold absolute concentrations are both defined by a standard deviation.

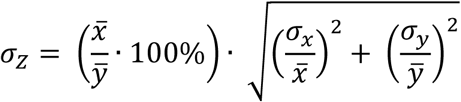

## Supporting information

Supporting information is available from the Wiley Online Library or from the authors.

## Acknowledgements

The authors thank Ulrike Hofmann for technical assistance and Matteo Castronovo for discussions. This work has received funding from the European Union’s EIC Pathfinder Challenges 2022 programme under grant agreement No 101115317 (NEO).

## Conflict of interest

The authors declare no conflict of interest.

## Data Availability Statement

The data that support the findings of this study are available in one of the Supplementary Files.

## Notes

### Competing Interest Statement

The authors have declared no competing interest.

### Summary of Updates

Additional quantification of ligation reactions in the presence of DMSO; additional control experiments; extended discussion; updated Experimental Section; updated author list

