## Supplementary information for "Global Quantitative Analysis of Ligation Reactions in Self-Assembled DNA Nanostructures at the Single-Nick Level"

1. B CUBE — Center for Molecular Bioengineering, Technische Universität Dresden, Dresden (Germany)
2. Technical and Macromolecular Chemistry, Paderborn University, Paderborn (Germany)
3. Dipartimento Interateneo di Fisica, Università degli Studi di Bari and INFN, Sezione di Bari, Bari (Italy)
4. These authors contribute equally to this work.

---

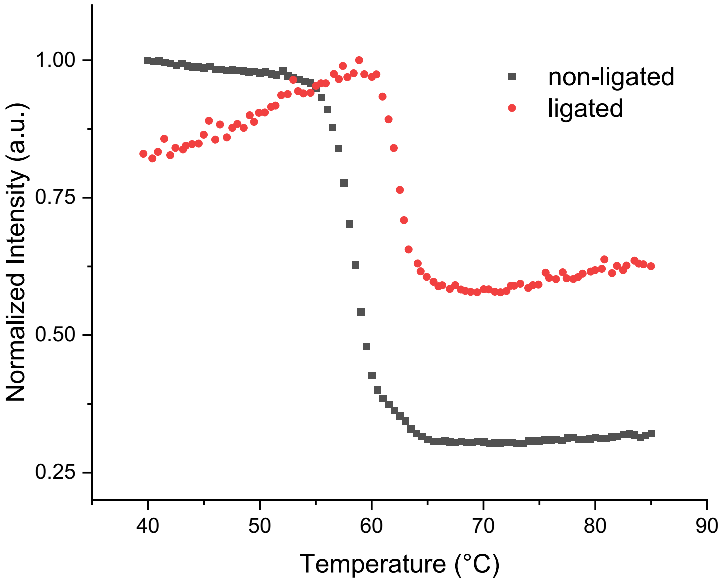


**Figure S1.** Normalized scattering intensity of ligated and non-ligated DON in water over a temperature range from 40°C to 85°C.

**Table 1**. *General thermal cycler program for quantification using qPCR*. Annealing temperature is dependent on the specific primer pair melting temperature*.

| steps | scan | temperature (°C) | time (m:s) | go to | loops | temperature increments (°C/s) |
|---|---|---|---|---|---|---|
| 1 | | 95 | 01:00 | | | 8 |
| 2 | | 95 | 00:10 | | | 6 |
| 3 | | ~50* | 00:05 | | | 6 |
| 4 | ♦ | 72 | 00:05 | 2 | 40 | 6 |
| 5 | | 60 | 00:05 | | | 6 |
| 6 | | 60 to 95 | 08:45 | | | $^{1}/_{15}$ |

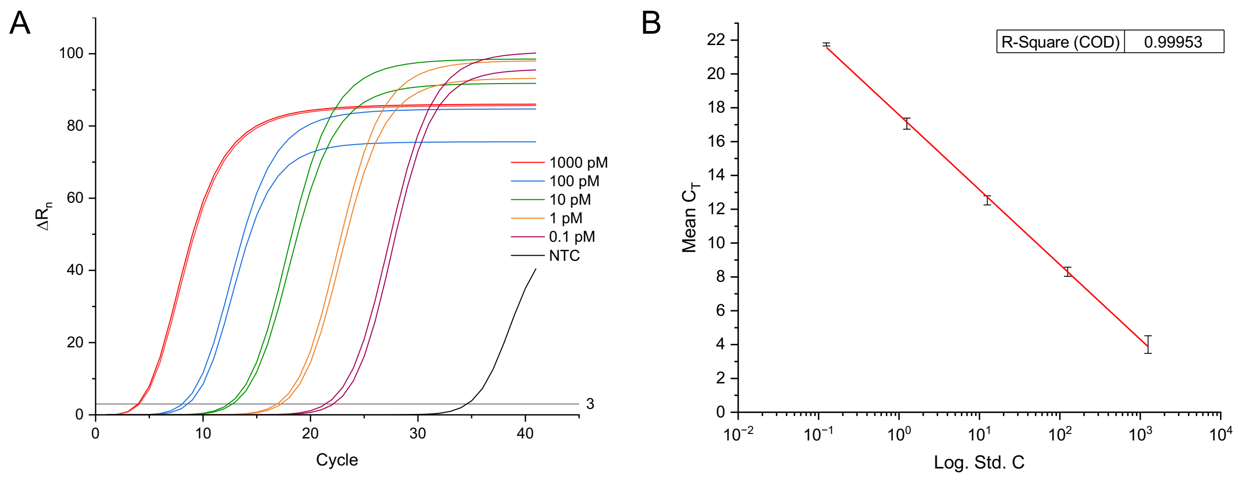


**Figure S2.** (A) Amplification curves of a selected sequence from the scaffold in tenfold dilutions and negative control (NTC). (B) Calibration curve of the selected sequence.

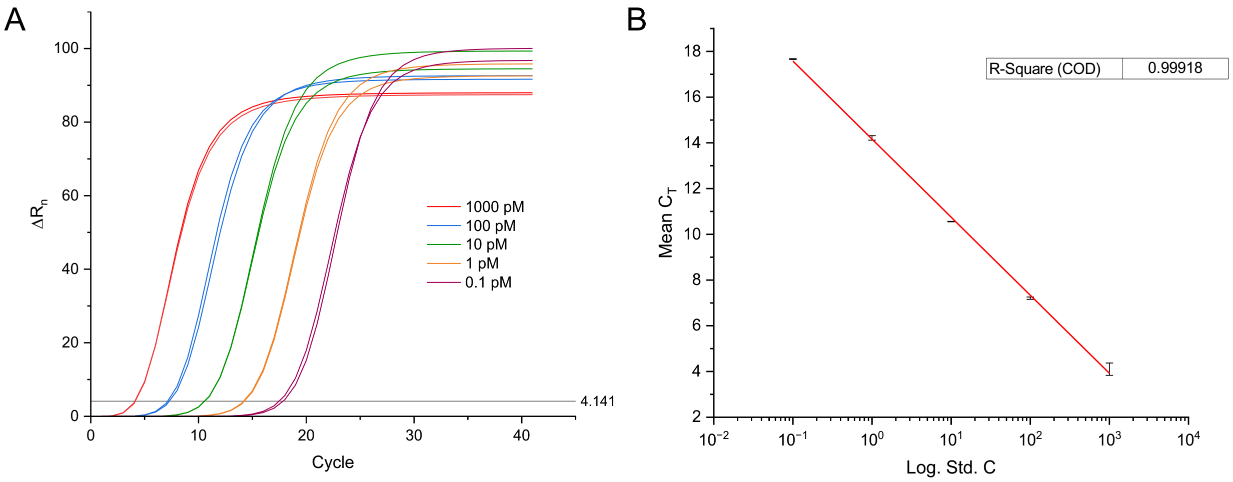


**Figure S3.** (A) Amplification curves of the sequence corresponding to the ligation product of the pair of staple strands at nick 1 in tenfold dilutions. (B) Calibration curve of the DNA sequence.

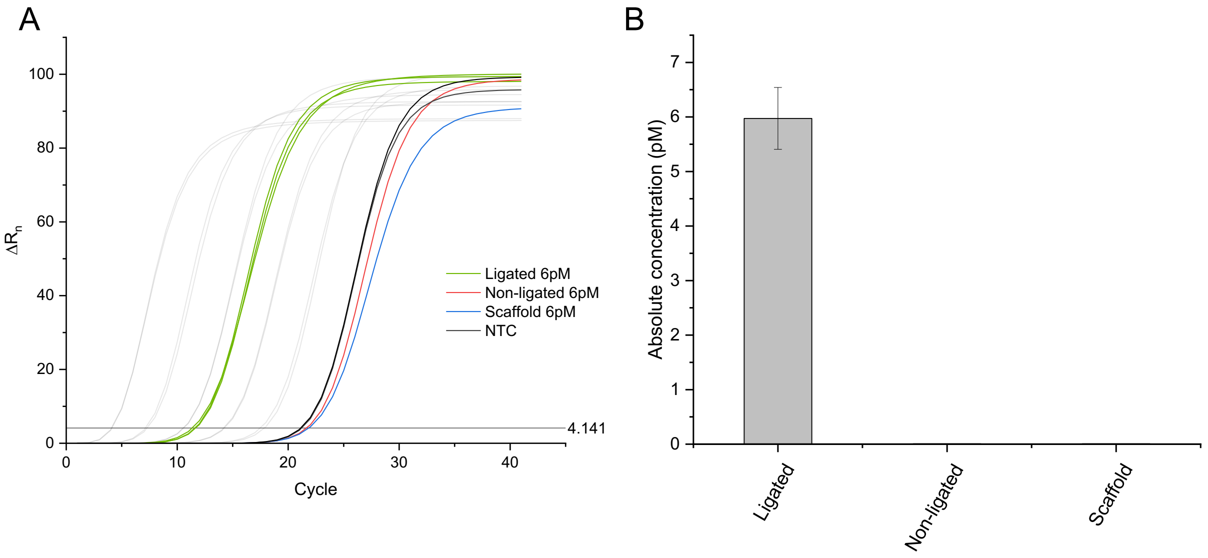


**Figure S4.** S*pecific amplification and quantification of the ligation product at nick 1 on DON, but not the staple strands and scaffold.* (A) Amplification curves of the ligated and non-ligated DON samples as well as the scaffold sample. (B) Calculated concentrations of the sequence with ligated, non-ligated and scaffold sample measured by qPCR and calculated according to the calibration curve in figure **S2B**.

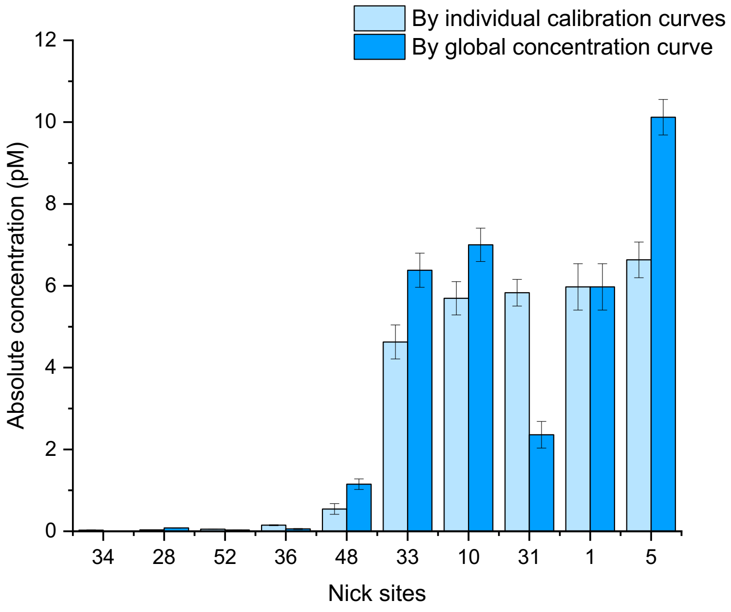


**Figure S5.** Comparison of the calculated concentrations derived from individual calibration curves or from the global calibration curve at nick 1 (figure **S2B**) for ten staple strand pairs.

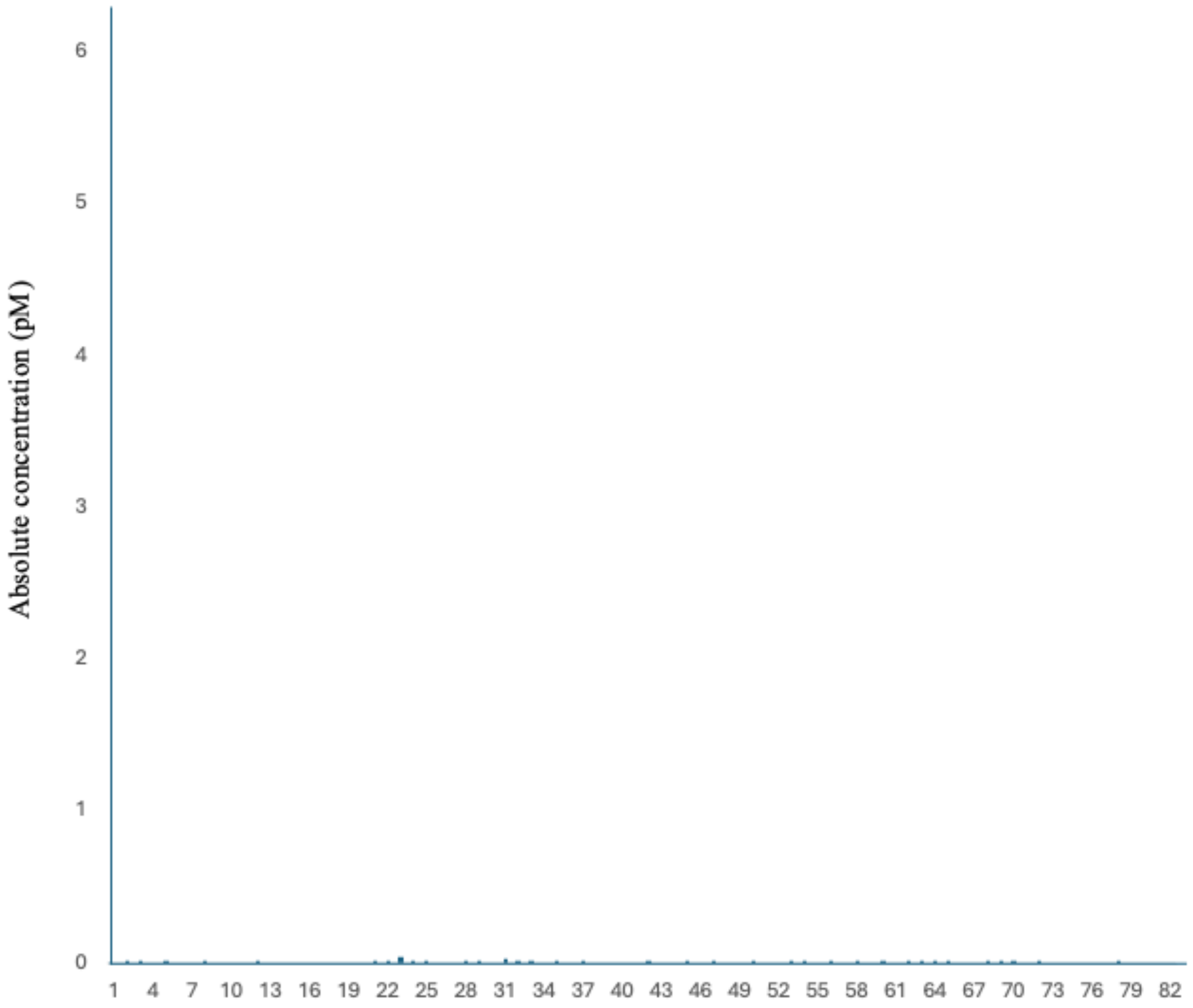


**Figure S6.** Shot gun analyses with 82 random pairs of primers**.**

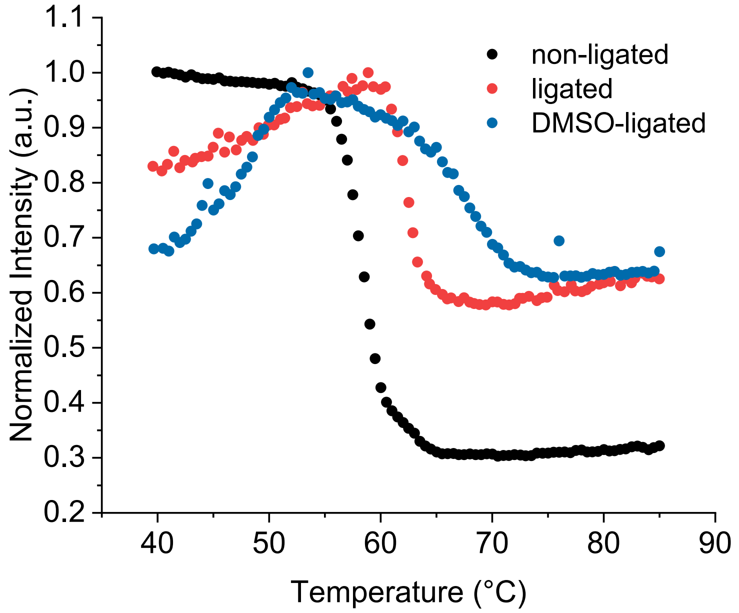


**Figure S7.** Temperature dependence of scattering intensity of triangular DNA origami structures without ligation, with ligation by T4 DNA ligase and with enzymatic ligation in the presence of DMSO.

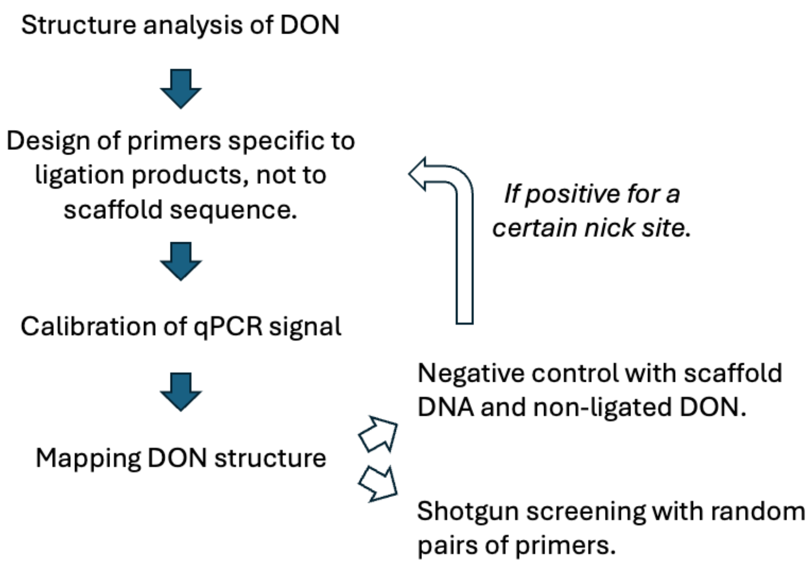


**Scheme S1**. Global quantitative analysis of ligation reactions work flow.

**Table S2.** *Staple strand pairs for ligation.*

| Nick # | Staple 1 | Sequence | Staple 2 | Sequence | Total sequence | Total length | Primer 1 | $T_m$ (°C) | Primer 2 | $T_m$ (°C) | $T_{anneal}$ (°C) |
|---|---|---|---|---|---|---|---|---|---|---|---|
| 1 | t-10s7h | ACGACAATAAATCCCGACTTGCGGGAGATCCTGAATCTTACCA | t-9s10g | ACGCTAACGAGCGTCTGGCGTTTTAGCGAACCCAACATGT | ACGACAATAAATCCCGACTTGCGGGAGATCCTGAATCTTACCAACGCTAACGAGCGTCTGGCGTTTTAGCGAACCCAACATGT | 83 | ACGACAATAAATCCCGACTT | 52 | ACATGTTGGGTTCGCTAAAA | 53 | 49 |
| 2 | t-7s8g | GCGCCTGTTATTCTAAGAACGCGATTCCAGAGCCTAATTT | t-7s10g | GCCAGTTACAAAATAATAGAAGGCTTATCCGGTTATCAAC | GCGCCTGTTATTCTAAGAACGCGATTCCAGAGCCTAATTTGCCAGTTACAAAATAATAGAAGGCTTATCCGGTTATCAAC | 80 | GCGCCTGTTATTCTAAGAAC | 53 | GTTGATAACCGGATAAGCCT | 53 | 49 |
| 3 | t-5s8g | ACAAGAAAGCAAGCAAATCAGATAACAGCCATATTATTTA | t-5s10g | TCCCAATCCAAATAAGATTACCGCGCCCAATAAATAATAT | ACAAGAAAGCAAGCAAATCAGATAACAGCCATATTATTTATCCCAATCCAAATAAGATTACCGCGCCCAATAAATAATAT | 72 | ACAAGAAAGCAAGCAAATCA | 57 | ATATTATTTATTGGGCGCGG | 51 | 49 |
| 4 | t-3s8g | AGCATGTATTTCATCGTAGGAATCAAACGATTTTTTGTTT | t-3s10g | AACGTCAAAAATGAAAAGCAAGCCGTTTTTATGAAACCAA | AGCATGTATTTCATCGTAGGAATCAAACGATTTTTTGTTTAACGTCAAAAATGAAAAGCAAGCCGTTTTTATGAAACCAA | 80 | AGCATGTATTTCATCGTAGGAATCA | 51 | TTGGTTTCATAAAAACGGCTTGCTT | 51 | 49 |
| 5 | t-1s8g | TTTCCTTAGCACTCATCGAGAACAATAGCAGCCTTTACAG | t-1s10e | AGAGAATAACATAAAAACAGGGAAGCGCATTA | TTTCCTTAGCACTCATCGAGAACAATAGCAGCCTTTACAGAGAGAATAACATAAAAACAGGGAAGCGCATTA | 72 | TTTCCTTAGCACTCATCGAGAACA | 57 | TAATGCGCTTCCCTGTTTTT | 53 | 50 |
| 6 | t-1s10e | AGAGAATAACATAAAAACAGGGAAGCGCATTA | t1s10g | GACGGGAGAATTAACTCGGAATAAGTTTATTTCCAGCGCC | AGAGAATAACATAAAAACAGGGAAGCGCATTAGACGGGAGAATTAACTCGGAATAAGTTTATTTCCAGCGCC | 72 | AGAGAATAACATAAAAACAGGGAAG | 51 | GGCGCTGGAAATAAACTTAT | 52 | 49 |
| 7 | t3s8g | CATTCAACAAACGCAAAGACACCAGAACACCCTGAACAAA | t3s10g | GTCAGAGGGTAATTGATGGCAACATATAAAAGCGATTGAG | CATTCAACAAACGCAAAGACACCAGAACACCCTGAACAAAGTCAGAGGGTAATTGATGGCAACATATAAAAGCGATTGAG | 80 | CATTCAACAAACGCAAAGAC | 52 | CTCAATCGCTTTTATATGTTGC | 51 | 49 |
| 8 | t5s8g | TTGACGGAAATACATACATAAAGGGCGCTAATATCAGAGA | t5s10g | GATAACCCACAAGAATGTTAGCAAACGTAGAAAATTATTC | TTGACGGAAATACATACATAAAGGGCGCTAATATCAGAGAGATAACCCACAAGAATGTTAGCAAACGTAGAAAATTATTC | 80 | TTGACGGAAATACATACATAAAG | 49 | GAATAATTTTCTACGTTTGCTAAC | 49 | 49 |
| 9 | t7s8g | CACCGTCACCTTATTACGCAGTATTGAGTTAAGCCCAATA | t7s10g | ATAAGAGCAAGAAACATGGCATGATTAAGACTCCGACTTG | CACCGTCACCTTATTACGCAGTATTGAGTTAAGCCCAATAATAAGAGCAAGAAACATGGCATGATTAAGACTCCGACTTG | 80 | CACCGTCACCTTATTACGCA | 57 | CAAGTCGGAGTCTTAATCAT | 50 | 49 |
| 10 | t9s8g | GAGCCAGCGAATACCCAAAAGAACATGAAATAGCAATAGC | t9s10h | TATCTTACCGAAGCCCAAACGCAATAATAACGAAAATCACCAG | GAGCCAGCGAATACCCAAAAGAACATGAAATAGCAATAGCTATCTTACCGAAGCCCAAACGCAATAATAACGAAAATCACCAG | 83 | GAGCCAGCGAATACCCAAAA | 57 | CTGGTGATTTTCGTTATTATTGC | 51 | 48 |

| Nick # | Staple 1 | Sequence | Staple 2 | Sequence | Total sequence | Total length | Primer 1 | $T_m$ (°C) | Primer 2 | $T_m$ (°C) | $T_{anneal}$ (°C) |
|---|---|---|---|---|---|---|---|---|---|---|---|
| 11 | t-12s9h | TGCTATTTTGCACCCAGCTACAATTTTGTTTTGAAGCCTTAAA | t-11s8e-t12s29c-0T | TCAAGATTAGTGTAGCAATACT | TGCTATTTTGCACCCAGCTACAATTTTGTTTTGAAGCCTTAAATCAAGATTAGTGTAGCAATACT | 65 | TGCTATTTTGCACCCAGCTA | 56 | AGTATTGCTACACTAATCTTGA | 49 | 46 |
| 12 | t1s8i | ATGGTTTATGTCACAATCAATAGATATTAAAC | t-1s8i | CAAGTACCTCATTCCAAGAACGGGAAATTCAT | ATGGTTTATGTCACAATCAATAGATATTAAACCAAGTACCTCATTCCAAGAACGGGAAATTCAT | 64 | ATGGTTTATGTCACAATCAATAGAT | 51 | ATGAATTTCCCGTTCTTGGA | 53 | 49 |
| 13 | t-11s18e-t12s9c-0T | ATAAGGCTTGCAACAAAGTTAC | t11s8h | CAGAAGGAAACCGAGGTTTTTAAGAAAAGTAAGCAGATAGCCG | ATAAGGCTTGCAACAAAGTTACCAGAAGGAAACCGAGGTTTTTAAGAAAAGTAAGCAGATAGCCG | 65 | ATAAGGCTTGCAACAAAGTT | 51 | CGGCTATCTGCTTACTTTTC | 52 | 49 |
| 14 | t-9s10g | ACGCTAACGAGCGTCTGGCGTTTTAGCGAACCCAACATGT | t-8s7c | TCAGCTAAAAAAGGTAAAGTAATT | ACGCTAACGAGCGTCTGGCGTTTTAGCGAACCCAACATGTTCAGCTAAAAAAGGTAAAGTAATT | 64 | ACGCTAACGAGCGTCTGGCG | 65 | AATTACTTTACCTTTTTTAGCTGA | 48 | 46 |
| 15 | t-8s5f | TTCTGACCTAAAATATAAAGTACCGACTGCAGAAC | t-7s8g | GCGCCTGTTATTCTAAGAACGCGATTCCAGAGCCTAATTT | TTCTGACCTAAAATATAAAGTACCGACTGCAGAACGCGCCTGTTATTCTAAGAACGCGATTCCAGAGCCTAATTT | 75 | TTCTGACCTAAAATATAAAGTACCG | 51 | AAATTAGGCTCTGGAATCGC | 54 | 49 |
| 16 | t-7s10g | GCCAGTTACAAAATAATAGAAGGCTTATCCGGTTATCAAC | t-6s7f | AATAGATAGAGCCAGTAATAAGAGATTTAATG | GCCAGTTACAAAATAATAGAAGGCTTATCCGGTTATCAACAATAGATAGAGCCAGTAATAAGAGATTTAATG | 80 | GCCAGTTACAAAATAATAGAAGG | 53 | CATTAAATCTCTTATTACTGGCT | 53 | 49 |
| 17 | t-5s6e | GTGTGATAAGGCAGAGGCATTTTCAGTCCTGA | t-5s8g | ACAAGAAAGCAAGCAAATCAGATAACAGCCATATTATTTA | GTGTGATAAGGCAGAGGCATTTTCAGTCCTGAACAAGAAAGCAAGCAAATCAGATAACAGCCATATTATTTA | 67 | GTGTGATAAGGCAGAGGCAT | 54 | TAAATAATATGGCTGTTATCTGATT | 55 | 49 |
| 18 | t-5s10g | TCCCAATCCAAATAAGATTACCGCGCCCAATAAATAATAT | t-4s7f | CCCATCCTCGCCAACATGTAATTTAATAAGGC | TCCCAATCCAAATAAGATTACCGCGCCCAATAAATAATATCCCATCCTCGCCAACATGTAATTTAATAAGGC | 80 | TCCCAATCCAAATAAGATTACC | 51 | GCCTTATTAAATTACATGTTGGC | 52 | 49 |
| 19 | t-3s6e | CACCGGAATCGCCATATTTAACAAAATTTACG | t-3s8g | AGCATGTATTTCATCGTAGGAATCAAACGATTTTTTGTTT | CACCGGAATCGCCATATTTAACAAAATTTACGAGCATGTATTTCATCGTAGGAATCAAACGATTTTTTGTTT | 72 | CACCGGAATCGCCATATTTAACAAA | 54 | AAACAAAAAATCGTTTGATTCCTA | 50 | 49 |
| 20 | t-3s10g | AACGTCAAAAATGAAAAGCAAGCCGTTTTTATGAAACCAA | t-2s7f | TCAATAATAGGGCTTAATTGAGAATCATAATT | AACGTCAAAAATGAAAAGCAAGCCGTTTTTATGAAACCAATCAATAATAGGGCTTAATTGAGAATCATAATT | 72 | AACGTCAAAAATGAAAAGCAA | 50 | AATTATGATTCTCAATTAAGCCC | 49 | 46 |

| Nick # | Staple 1 | Sequence | Staple 2 | Sequence | Total sequence | Total length | Primer 1 | $T_m$ (°C) | Primer 2 | $T_m$ (°C) | $T_{anneal}$ (°C) |
|---|---|---|---|---|---|---|---|---|---|---|---|
| 21 | t-1s6e | TTAGTATCGCCAACGCTCAACAGTCGGCTGTC | t-1s8g | TTTCCTTAGCACTCATCGAGAACAATAGCAGCCTTTACAG | TTAGTATCGCCAACGCTCAACAGTCGGCTGTCTTTCCTTAGCACTCATCGAGAACAATAGCAGCCTTTACAG | 72 | TTAGTATCGCCAACGCTCAA | 55 | CTGTAAAGGCTGCTATTGTT | 51 | 49 |
| 22 | t-1s8i | CAAGTACCTCATTCCAAGAACGGGAAATTCAT | t1s8i | ATGGTTTATGTCACAATCAATAGATATTAAAC | CAAGTACCTCATTCCAAGAACGGGAAATTCATATGGTTTATGTCACAATCAATAGATATTAAAC | 64 | CAAGTACCTCATTCCAAGAA | 50 | GTTTAATATCTATTGATTGTGACAT | 47 | 46 |
| 23 | t1s10g | GACGGGAGAATTAACTCGGAATAAGTTTATTTCCAGCGCC | t2s7f | AAAGACAACATTTTCGGTCATAGCCAAAATCA | GACGGGAGAATTAACTCGGAATAAGTTTATTTCCAGCGCCAAAGACAACATTTTCGGTCATAGCCAAAATCA | 72 | GACGGGAGAATTAACTCGGA | 56 | TGATTTTGGCTATGACCGAA | 53 | 49 |
| 24 | t3s6e | CACCGGAAAGCGCGTTTTCATCGGAAGGGCGA | t3s8g | CATTCAACAAACGCAAAGACACCAGAACACCCTGAACAAA | CACCGGAAAGCGCGTTTTCATCGGAAGGGCGACATTCAACAAACGCAAAGACACCAGAACACCCTGAACAAA | 72 | CACCGGAAAGCGCGTTTTCA | 61 | TTTGTTCAGGGTGTTCTGGT | 56 | 54 |
| 25 | t3s10g | GTCAGAGGGTAATTGATGGCAACATATAAAAGCGATTGAG | t4s7f | GGAGGGAATTTAGCGTCAGACTGTCCGCCTCC | GTCAGAGGGTAATTGATGGCAACATATAAAAGCGATTGAGGGAGGGAATTTAGCGTCAGACTGTCCGCCTCC | 72 | GTCAGAGGGTAATTGATGGC | 55 | GGAGGCGGACAGTCTGACGC | 66 | 54 |
| 26 | t5s6e | TCAGAACCCAGAATCAAGTTTGCCGGTAAATA | t5s8g | TTGACGGAAATACATACATAAAGGGCGCTAATATCAGAGA | TCAGAACCCAGAATCAAGTTTGCCGGTAAATATTGACGGAAATACATACATAAAGGGCGCTAATATCAGAGA | 72 | TCAGAACCCAGAATCAAGTT | 52 | TCTCTGATATTAGCGCCCTT | 54 | 49 |
| 27 | t5s10g | GATAACCCACAAGAATGTTAGCAAACGTAGAAAATTATTC | t6s7f | ATTAAAGGCCGTAATCAGTAGCGAGCCACCCT | GATAACCCACAAGAATGTTAGCAAACGTAGAAAATTATTCATTAAAGGCCGTAATCAGTAGCGAGCCACCCT | 72 | GATAACCCACAAGAATGTTAGC | 53 | AGGGTGGCTCGCTACTGATT | 61 | 49 |
| 28 | t7s6e | AGAGCCGCACCATCGATAGCAGCATGAATTAT | t7s8g | CACCGTCACCTTATTACGCAGTATTGAGTTAAGCCCAATA | AGAGCCGCACCATCGATAGCAGCATGAATTATCACCGTCACCTTATTACGCAGTATTGAGTTAAGCCCAATA | 72 | AGAGCCGCACCATCGATAGC | 62 | TATTGGGCTTAACTCAATACTGC | 54 | 49 |
| 29 | t7s10g | ATAAGAGCAAGAAACATGGCATGATTAAGACTCCGACTTG | t8s7g | AGCCATTTAAACGTCACCAATGAACACCAGAACCA | ATAAGAGCAAGAAACATGGCATGATTAAGACTCCGACTTGAGCCATTTAAACGTCACCAATGAACACCAGAACCA | 75 | ATAAGAGCAAGAAACATGGC | 52 | TGGTTCTGGTGTTCATTGGT | 56 | 49 |
| 30 | t9s6e | CCATTAGCAAGGCCGGGGGAATTA | t9s8g | GAGCCAGCGAATACCCAAAAGAACATGAAATAGCAATAGC | CCATTAGCAAGGCCGGGGGAATTAGAGCCAGCGAATACCCAAAAGAACATGAAATAGCAATAGC | 64 | CCATTAGCAAGGCCGGGGGA | 65 | GCTATTGCTATTTCATGTTCTTT | 50 | 49 |

| Nick # | Staple 1 | Sequence | Staple 2 | Sequence | Total sequence | Total length | Primer 1 | $T_m$ (°C) | Primer 2 | $T_m$ (°C) | $T_{anneal}$ (°C) |
|---|---|---|---|---|---|---|---|---|---|---|---|
| 31 | t-8s7c | TCAGCTAAAAAAGGTAAAGTAATT | t-9s6e-t10s27c-1T | TCAGCTAAAAAAGGTAAAGTAATT | TCAGCTAAAAAAGGTAAAGTAATTCTGTCCAGACGTATACCGAACGA | 47 | TCAGCTAAAAAAGGTAAAGTAATT | 48 | TCGTTCGGTATACGTCTGGA | 56 | 46 |
| 32 | t1s6i | TTCATAATCCCCTTATTAGCGTTTTTCTTACC | t-1s6i | AGTATAAAATATGCGTTATACAAAGCCATCTT | TTCATAATCCCCTTATTAGCGTTTTTCTTACCAGTATAAAATATGCGTTATACAAAGCCATCTT | 64 | TTCATAATCCCCTTATTAGCG | 50 | AAGATGGCTTTGTATAACGC | 52 | 49 |
| 33 | t-9s16e-t10s7c-1T | ACTACGAAGGCTTAGCACCATTA | t9s6e | ACTACGAAGGCTTAGCACCATTA | ACTACGAAGGCTTAGCACCATTACCATTAGCAAGGCCGGGGGAATTA | 47 | ACTACGAAGGCTTAGCACCA | 57 | TAATTCCCCCGGCCTTGCTA | 60 | 54 |
| 34 | t-6s7f | AATAGATAGAGCCAGTAATAAGAGATTTAATG | t-6s5c | GTTTGAAATTCAAATATATTTTAG | AATAGATAGAGCCAGTAATAAGAGATTTAATGGTTTGAAATTCAAATATATTTTAG | 56 | AATAGATAGAGCCAGTAATA | 44 | CTAAAATATATTTGAATTTCAAAC | 43 | 40 |
| 35 | t-6s3f | TCCCTTAGAATAACGCGAGAAAACTTTTACCGACC | t-5s6e | GTGTGATAAGGCAGAGGCATTTTCAGTCCTGA | TCCCTTAGAATAACGCGAGAAAACTTTTACCGACCGTGTGATAAGGCAGAGGCATTTTCAGTCCTGA | 72 | TCCCTTAGAATAACGCGAGA | 50 | TCAGGACTGAAAATGCCTCT | 48 | 46 |
| 36 | t-4s7f | CCCATCCTCGCCAACATGTAATTTAATAAGGC | t-4s5f | GTTAAATACAATCGCAAGACAAAGCCTTGAAA | CCCATCCTCGCCAACATGTAATTTAATAAGGCGTTAAATACAATCGCAAGACAAAGCCTTGAAA | 64 | CCCATCCTCGCCAACATGTA | 59 | TTTCAAGGCTTTGTCTTGCG | 55 | 52 |
| 37 | t-3s4e | GATTAAGAAATGCTGATGCAAATCAGAATAAA | t-3s6e | CACCGGAATCGCCATATTTAACAAAATTTACG | GATTAAGAAATGCTGATGCAAATCAGAATAAACACCGGAATCGCCATATTTAACAAAATTTACG | 64 | GATTAAGAAATGCTGATGCAA | 50 | CGTAAATTTTGTTAAATATGGCGA | 50 | 49 |
| 38 | t-2s7f | TCAATAATAGGGCTTAATTGAGAATCATAATT | t-2s5f | ACTAGAAATATATAACTATATGTACGCTGAGA | TCAATAATAGGGCTTAATTGAGAATCATAATTACTAGAAATATATAACTATATGTACGCTGAGA | 64 | TCAATAATAGGGCTTAATTGAGA | 49 | TCTCAGCGTACATATAGTTA | 48 | 46 |
| 39 | t-1s4e | TTATCAAACCGGCTTAGGTTGGGTAAGCCTGT | t-1s6e | TTAGTATCGCCAACGCTCAACAGTCGGCTGTC | TTATCAAACCGGCTTAGGTTGGGTAAGCCTGTTTAGTATCGCCAACGCTCAACAGTCGGCTGTC | 64 | TTATCAAACCGGCTTAGGTT | 53 | GACAGCCGACTGTTGAGCGT | 63 | 49 |
| 40 | t-1s6i | AGTATAAAATATGCGTTATACAAAGCCATCTT | t1s6i | TTCATAATCCCCTTATTAGCGTTTTTCTTACC | AGTATAAAATATGCGTTATACAAAGCCATCTTTTCATAATCCCCTTATTAGCGTTTTTCTTACC | 64 | AGTATAAAATATGCGTTATACAAAG | 47 | GGTAAGAAAAACGCTAATAAG | 47 | 46 |

| Nick # | Staple 1 | Sequence | Staple 2 | Sequence | Total sequence | Total length | Primer 1 | $T_m$ (°C) | Primer 2 | $T_m$ (°C) | $T_{anneal}$ (°C) |
|---|---|---|---|---|---|---|---|---|---|---|---|
| 41 | t2s7f | AAAGACAACATTTTCGGTCATAGCCAAAATCA | t2s5f | CCGGAACCCAGAATGGAAAGCGCAACATGGCT | AAAGACAACATTTTCGGTCATAGCCAAAATCACCGGAACCCAGAATGGAAAGCGCAACATGGCT | 64 | AAAGACAACATTTTCGGTCA | 51 | AGCCATGTTGCGCTTTCCAT | 60 | 49 |
| 42 | **t3s4e** | **TGTACTGGAAATCCTCATTAAAGCAGAGCCAC** | **t3s6e** | **CACCGGAAAGCGCGTTTTCATCGGAAGGGCGA** | **TGTACTGGAAATCCTCATTAAAGCAGAGCCACCACCGGAAAGCGCGTTTTCATCGGAAGGGCGA** | **64** | **TGTACTGGAAATCCTCATTAAA** | **50** | **TCGCCCTTCCGATGAAAACG** | **60** | **49** |
| 43 | t4s7f | GGAGGGAATTTAGCGTCAGACTGTCCGCCTCC | t4s5f | CTCAGAGCATATTCACAAACAAATTAATAAGT | GGAGGGAATTTAGCGTCAGACTGTCCGCCTCCCTCAGAGCATATTCACAAACAAATTAATAAGT | 64 | GGAGGGAATTTAGCGTCAGA | 56 | ACTTATTAATTTGTTTGTGAATATG | 46 | 41 |
| 44 | **t5s4e** | **CCTTGAGTCAGACGATTGGCCTTGCGCCACCC** | **t5s6e** | **TCAGAACCCAGAATCAAGTTTGCCGGTAAATA** | **CCTTGAGTCAGACGATTGGCCTTGCGCCACCCTCAGAACCCAGAATCAAGTTTGCCGGTAAATA** | **64** | **CCTTGAGTCAGACGATTGGC** | **58** | **TATTTACCGGCAAACTTGAT** | **50** | **49** |
| 45 | **t6s7f** | **ATTAAAGGCCGTAATCAGTAGCGAGCCACCCT** | **t6s5g** | **CAGAGCCAGGAGGTTGAGGCAGGTAACAGTGCCCG** | **ATTAAAGGCCGTAATCAGTAGCGAGCCACCCTCAGAGCCAGGAGGTTGAGGCAGGTAACAGTGCCCG** | **67** | **ATTAAAGGCCGTAATCAGTAG** | **51** | **CGGGCACTGTTACCTGCCTC** | **64** | **49** |
| 46 | **t7s4e** | **GCCGCCAGCATTGACACCACCCTC** | **t7s6e** | **AGAGCCGCACCATCGATAGCAGCATGAATTAT** | **GCCGCCAGCATTGACACCACCCTCAGAGCCGCACCATCGATAGCAGCATGAATTAT** | **56** | **GCCGCCAGCATTGACACCAC** | **65** | **ATAATTCATGCTGCTATCGATG** | **51** | **48** |
| 47 | **t-6s5c** | **GTTTGAAATTCAAATATATTTTAG** | **t-7s4e-t8s25c-2T** | **TTAATTTCATCTTAGACTTTACAA** | **GTTTGAAATTCAAATATATTTTAGTTAATTTCATCTTAGACTTTACAA** | **48** | **GTTTGAAATTCAAATATATTTTAG** | **42** | **TTGTAAAGTCTAAGATGAAA** | **44** | **41** |
| 48 | **t1s4i** | **AGCGTCATGTCTCTGAATTTACCGACTACCTT** | **t-1s4i** | **TTTAACCTATCATAGGTCTGAGAGTTCCAGTA** | **AGCGTCATGTCTCTGAATTTACCGACTACCTTTTTAACCTATCATAGGTCTGAGAGTTCCAGTA** | **64** | **AGCGTCATGTCTCTGAATTT** | **52** | **TACTGGAACTCTCAGACCTA** | **52** | **49** |
| 49 | **t-7s14e-t8s5c-2T** | **ATGACCCTGTAATACTTCAGAGCA** | **t7s4e** | **GCCGCCAGCATTGACACCACCCTC** | **ATGACCCTGTAATACTTCAGAGCAGCCGCCAGCATTGACACCACCCTC** | **48** | **TTTAATTGTATTTCCACCAGAG** | **49** | **GAGGGTGGTGTCAATGCTGG** | **61** | **46** |
| 50 | **t-4s5f** | **GTTAAATACAATCGCAAGACAAAGCCTTGAAA** | **t-4s3g** | **ACATAGCGCTGTAAATCGTCGCTATTCATTTCAATTACCT** | **GTTAAATACAATCGCAAGACAAAGCCTTGAAAACATAGCGCTGTAAATCGTCGCTATTCATTTCAATTACCT** | **72** | **GTTAAATACAATCGCAAGACAAAG** | **51** | **AGGTAATTGAAATGAATAGCGAC** | **51** | **49** |

| Nick # | Staple 1 | Sequence | Staple 2 | Sequence | Total sequence | Total length | Primer 1 | $T_m$ (°C) | Primer 2 | $T_m$ (°C) | $T_{anneal}$ (°C) |
|---|---|---|---|---|---|---|---|---|---|---|---|
| 51 | t-4s1g | GAGCAAAAGAAGATG<br>AGTGAATAACCTTGCT<br>TATAGCTTA | t-3s4e | GATTAAGAAATGCTG<br>ATGCAAATCAGAATA<br>AA | GAGCAAAAGAAGATGAGTGAATAA<br>CCTTGCTTATAGCTTAGATTAAGAA<br>ATGCTGATGCAAATCAGAATAAA | 72 | GAGCAAAAGA<br>AGATGAGTGA | 51 | TTTATTCTGA<br>TTTGCATCA<br>GC | 50 | 49 |
| 52 | t-2s5f | ACTAGAAATATATAACT<br>ATATGTACGCTGAGA | t-2s1g | AAAACAAAATTAATT<br>AAATGGAAACAGTAC<br>ATTAGTGAAT | ACTAGAAATATATAACTATATGTACG<br>CTGAGAAAAACAAAATTAATTAAA<br>TGGAAACAGTACATTAGTGAAT | 72 | AGAGTCAAAA<br>ATCAATATATGT<br>GAT | 48 | ATTCACTAAT<br>GTACTGTTT<br>CC | 48 | 46 |
| 53 | t-2s1g | AAAACAAAATTAATTA<br>AATGGAAACAGTACAT<br>TAGTGAAT | t-1s4e | TTATCAAACCGGCTT<br>AGGTTGGGTAAGCCT<br>GT | AAAACAAAATTAATTAAATGGAAA<br>CAGTACATTAGTGAATTTATCAAAC<br>CGGCTTAGGTTGGGTAAGCCTGT | 72 | AAAACAAAATT<br>AATTAAATGGA<br>AAC | 45 | ACAGGCTTA<br>CCCAACCTA<br>AG | 56 | 41 |
| 54 | t-1s4i | TTTAACCTATCATAGGT<br>CTGAGAGTTCCAGTA | t1s4i | AGCGTCATGTCTCTG<br>AATTTACCGACTACC<br>TT | TTTAACCTATCATAGGTCTGAGAGT<br>TCCAGTAAGCGTCATGTCTCTGAAT<br>TTACCGACTACCTT | 64 | TTTAACCTATCA<br>TAGGTCTGAG | 49 | AAGGTAGTC<br>GGTAAATTC<br>AG | 50 | 46 |
| 55 | t2s5f | CCGGAACCCAGAATG<br>GAAAGCGCAACATGG<br>CT | t2s3g | TTTGATGATTAAGAG<br>GCTGAGACTTGCTCA<br>GTACCAGGCG | CCGGAACCCAGAATGGAAAGCGC<br>AACATGGCTTTTGATGATTAAGAGG<br>CTGAGACTTGCTCAGTACCAGGCG | 72 | CCGGAACCCAG<br>AATGGAAAG | 58 | CGCCTGGTA<br>CTGAGCAAG<br>TC | 61 | 54 |
| 56 | t2s1g | GATAAGTGCCGTCGAG<br>CTGAAACATGAAAGTA<br>TACAGGAG | t3s4e | TGTACTGGAAATCCT<br>CATTAAAGCAGAGCC<br>AC | GATAAGTGCCGTCGAGCTGAAACA<br>TGAAAGTATACAGGAGTGTACTGG<br>AAATCCTCATTAAAGCAGAGCCAC | 72 | GATAAGTGCCG<br>TCGAGCTGA | 59 | GTGGCTCTG<br>CTTTAATGA<br>GG | 55 | 54 |
| 57 | t4s5f | CTCAGAGCATATTCAC<br>AAACAAATTAATAAGT | t4s3g | TTTAACGGTTCGGAA<br>CCTATTATTAGGGTTG<br>ATATAAGTA | CTCAGAGCATATTCACAAACAAATT<br>AATAAGTTTTAACGGTTCGGAACCT<br>ATTATTAGGGTTGATATAAGTA | 72 | CTCAGAGCATA<br>TTCACAAAC | 50 | TACTTATATC<br>AACCCTAAT<br>AATAGG | 47 | 46 |
| 58 | t4s1g | TAGCCCGGAATAGGTG<br>AATGCCCCCTGCCTAT<br>GGTCAGTG | t5s4e | CCTTGAGTCAGACGA<br>TTGGCCTTGCGCCAC<br>CC | TAGCCCGGAATAGGTGAATGCCCC<br>CTGCCTATGGTCAGTGCCTTGAGTC<br>AGACGATTGGCCTTGCGCCACCC | 72 | TAGCCCGGAAT<br>AGGTGAATG | 56 | GGGTGGCGC<br>AAGGCCAAT<br>CG | 68 | 54 |
| 59 | t1s2i | CGGGGTTTCCTCAAGA<br>GAAGGATTTTGAATTA | t-1s2i | CCTTTTTTCATTTAAC<br>AATTTCATAGGATTAG | CGGGGTTTCCTCAAGAGAAGGATT<br>TTGAATTACCTTTTTTCATTTAACA<br>ATTTCATAGGATTAG | 64 | CGGGGTTTCCT<br>CAAGAGAAG | 57 | CTAATCCTAT<br>GAAATTGTT<br>AAATGA | 47 | 46 |
| 60 | t-4s3g | ACATAGCGCTGTAAAT<br>CGTCGCTATTCATTTCA<br>ATTACCT | t-4s1g | GAGCAAAAGAAGAT<br>GAGTGAATAACCTTG<br>CTTATAGCTTA | ACATAGCGCTGTAAATCGTCGCTAT<br>TCATTTCAATTACCTGAGCAAAAG<br>AAGATGAGTGAATAACCTTGCTTAT<br>AGCTTA | 80 | ACATAGCGCTG<br>TAAATCGTC | 54 | TAAGCTATA<br>AGCAAGGTT<br>ATTCA | 50 | 49 |

| Nick # | Staple 1 | Sequence | Staple 2 | Sequence | Total sequence | Total length | Primer 1 | $T_m$ (°C) | Primer 2 | $T_m$ (°C) | $T_{anneal}$ (°C) |
|---|---|---|---|---|---|---|---|---|---|---|---|
| 61 | t-2s3g | AGAGTCAAAAATCAATATATGTGATGAAACAAACATCAAG | t-2s1g | AAAACAAAATTAATTAAATGGAAACAGTACATTAGTGAAT | AGAGTCAAAAATCAATATATGTGATGAAACAAACATCAAGAAAACAAAATTAATTAAATGGAAACAGTACATTAGTGAAT | 80 | AGAGTCAAAAATCAATATATGTGAT | 48 | ATTCACTAATGTACTGTTTCC | 48 | 46 |
| 62 | t-1s2i | CCTTTTTTCATTTAACAATTTCATAGGATTAG | t1s2i | CGGGGTTTCCTCAAGAGAAGGATTTTGAATTA | CCTTTTTTCATTTAACAATTTCATAGGATTAGCGGGGTTTCCTCAAGAGAAGGATTTTGAATTA | 64 | CCTTTTTTCATTTAACAATTTCATA | 46 | TAATTCAAAATCCTTCTCTTGA | 47 | 46 |
| 63 | t2s3g | TTTGATGATTAAGAGGCTGAGACTTGCTCAGTACCAGGCG | t2s1g | GATAAGTGCCGTCGAGCTGAAACATGAAAGTATACAGGAG | TTTGATGATTAAGAGGCTGAGACTTGCTCAGTACCAGGCGGATAAGTGCCGTCGAGCTGAAACATGAAAGTATACAGGAG | 80 | TTTGATGATTAAGAGGCTGA | 49 | CTCCTGTATACTTTCATGTT | 47 | 46 |
| 64 | t4s3g | TTTAACGGTTCGGAACCTATTATTAGGGTTGATATAAGTA | t4s1g | TAGCCCGGAATAGGTGAATGCCCCCTGCCTATGGTCAGTG | TTTAACGGTTCGGAACCTATTATTAGGGTTGATATAAGTATAGCCCGGAATAGGTGAATGCCCCCTGCCTATGGTCAGTG | 80 | TTTAACGGTTCGGAACCTAT | 52 | CACTGACCATAGGCAGGGGG | 63 | 49 |

**Table S3.** *Oligonucleotides for calibrations.*

| Experiment | Sequence | Length (nt) | Forward primer | $T_m$ (°C) | Reverse primer | $T_m$ (°C) | $T_{anneal}$ (°C) |
|---|---|---|---|---|---|---|---|
| Absolute quantification of DNA nanostructures' scaffold strand | CTGTTGCAGGCGGTGTTAATACTGA CCGCCTCACCTCTGTTTTATCTTCT GCTGGTGGTTCGTT | 64 | AACGAACCACCAGCAGAAGA | 58 | CTGTTGCAGGCGGTGTTAAT | 58 | 52 |
| Absolute quantification of ligated staple strands | ACGACAATAAATCCCGACTTGCGG GAGATCCTGAATCTTACCAACGCTA ACGAGCGTCTGGCGTTTTAGCGAA CCCAACATGT | 83 | ACGACAATAAATCCCGACTT | 52 | ACATGTTGGGTTCGCTAAAA | 53 | 49 |
